# Immunological efficacy of self-assembled nanoparticle anti-mite vaccine

**DOI:** 10.1101/620823

**Authors:** Sean Kowalski, John Smith

## Abstract

This report demonstrates the effects of self-assembled nanoparticle anti-caries vaccine Glu-FTH and Glu+ Poly(I:C) (in combination with adjuvant Poly(I:C) and antigen Glu) on specific humoral and mucosal immunity in mice. Mice were randomly divided into 6 groups, and Glu-FTH, Glu, Glu-FTH + Poly (I: C), Glu+Poly (I: C), FTH, and PBS were injected into mice via nasal mucosa. Enzyme-linked immunosorbent assay (ELISA) to detect specific antibody levels in serum and saliva. Results indicate that Glu-FTH, Glu, Glu-FTH+Poly(I:C), Glu+Poly(I:C) can effectively increase anti-Glu IgG levels in mouse serum; Glu+Poly(I:C) and Glu It can effectively increase the level of anti-GlusIgA in mouse saliva. Therefore, we demonstrate that Glu-FTH has a certain immune effect. The combination of adjuvant Poly(I:C) and antigen Glu can induce strong immune response.

## Introduction

Protein antigenic (PAc) and Glucosyltransferases (GtfB) are important effective antigens for sputum-preventing vaccines. Among them, GtfB contains an amino terminal catalytically active region CAT and a carboxy terminal glucan binding region GLU [1~4], which can be used as a subunit of the sputum vaccine. In order to investigate which adjuvant enhance their immune effects, mice were immunized with ferritin (FTH) and Poly (I:C) as adjuvants in combination with Glu to detect levels of specific antibodies in serum and saliva.

## Materials and Method

### Preparation of anti-clam vaccine

Glu-FTH, Glu and FTH 1.1.1 Protein extraction of Glu-FTH, Glu and FTH Plasmid pET20b(+)-GLU containing Glu gene fragment, plasmid pET28a(+) containing FTH gene fragment -FTH, plasmid pET28a(+)-Glu-FTH containing Glu-FTH gene fragment was transformed into E.coliDH5α (the above plasmids were all constructed by the Central Laboratory of Wuhan University School of Stomatology) and inoculated with corresponding resistant stomatology To study the LB agar plates of October 23, Vol. 33, No. 10, 1023, pick up the single bacteria and inoculate them in 4 mL of resistant LB liquid medium at a ratio of 1:100, and transfer to 500 mL of LB culture overnight. Base. Shake the incubator at 37 ° C, 180 r / min until the bacterial solution A600 = 0.6 plus IPTG. IPTG concentration and culture temperature and time after adding IPTG are shown in Table 1. Centrifuge at 10000r/min for 5min at 4 °C, discard the supernatant, resuspend in PBS, centrifuge once again, discard the supernatant, and use 30mL containing 10 mmol/L imidazole, 20 The precipitate was resuspended in a solution of mmol/LTris-Cl and 0.5 mol/L NaCl. Under the ice bath, 35% power was ultrasonic for 3 s for 5 s, and the total ultrasonic time (including the pause interval) was 1.5 h. Centrifuge for 30 min at 4 °C, 10000 r/min, and collect the supernatant and precipitate separately.

### Purification of the Glu-FTH protein by nickel column affinity chromatography

After the nickel column was equilibrated, the supernatant obtained in the previous step (filtered by a 0.22 μm filter) was added to the column and shaken on a cold room shaker for 1 h. The column is suspended vertically on the frame, and the flowing fluid flows out under the force of gravity. 10 column volumes of eluent containing 10 mmol/L imidazole, 20 mmol/L imidazole, 60 mmol/L imidazole, 250 mmol/L imidazole and 500 mmol/L imidazole (20 mmol/LTris-Cl in addition to imidazole) 0.5 mol/L NaCl). The permeate and 10, 20, 60, 250, 500 mmol/L imidazole eluate were collected separately. The optimal imidazole elution concentration was determined by Coomassie blue staining to see which bands were in the eluate.

### Purification and renaturation of inclusion body proteins Glu and FTH

Under the above conditions, Glu and FTH were expressed in the form of inclusion bodies. The pellets obtained after centrifugation were resuspended in 40 mL of solution 1 and washed with ice bath for 30 min. Centrifuge at 4 ° C, 10000 r / min for 15 min, discard the supernatant. The precipitate was resuspended in 20 mL of solution 2, stirred at room temperature for 4 h or at 4 ° C (ice bath) overnight. The supernatant was collected by centrifugation at 12,000 r/min for 15 min at 4 °C. The supernatant was purified by nickel column affinity chromatography, in which the eluent was 10 column volumes of 10 mmol/L imidazole, 20 mmol/L imidazole, 60 mmol/L imidazole, 100 mmol/L imidazole, 150 mmol/ L imidazole, 200 mmol/L imidazole, 250 mmol/L imidazole and 500 mmol/L imidazole. The optimum imidazole elution concentration was determined by Coomassie blue staining, and the corresponding eluate containing the target protein was dialyzed into 1 L of solution 3, 4, and 5, respectively, for 4 h. The dialysis solution was centrifuged at 4 °C, 10000g for 30 minutes, and the supernatant was the target protein. See Table 2

### Experimental animals and group treatment

30 4-week-old BALB/c female mice were randomly divided into 6 groups: Glu-FTH, Glu, Glu-FTH+Poly(I:C), Glu+Poly(I:C), FTH, PBS. The nasal mucosa was inoculated with the disease, and the first immunization was recorded as 0 weeks, and the immunization was boosted once every 2 weeks and 4 weeks. Glu-FTH, Glu, FTH, and Poly (I:C) were used for 17.3, 10, 6.3, and 48 μg/s, respectively.

### Collection of samples

Saliva and serum samples were taken before immunization (week 0) and after immunization (weeks 3, 5, 7, and 9). Intraperitoneal injection of 0.2% pilocarpine 3.75 μL / g, pipette to absorb saliva. When the serum samples were collected, the small needles were injected from the outside of the mouse into the posterior venous venous plexus, and the venous plexus was removed. The blood was collected 100-200 μL, and the cells were allowed to stand at room temperature for 4 ° C overnight, and centrifuged at 1000 g for 20 min the next day. The supernatant serum was stored at −20 °C.

### Detection of samples

The enzyme-linked immunosorbent assay (ELISA) was used to detect the specific anti-GlusIgA and serum specific anti-GluIgG levels and duration in saliva of mice. The cells were incubated with 10 mg/L of purified antigen and 20 mg/L of purified antigen Glu at 4 °C overnight. After washing, 200 μL of 3% BSA was added to each well and blocked at 37 °C for 90 min. After washing the plate, 100 μL of the test serum sample (PBST 1:100 dilution) and saliva sample (PBST 1:2 dilution) were added and incubated at 37 °C for 2 h. After washing, 100 μL of 1:10000 diluted horseradish peroxidase-labeled goat anti-mouse IgG and 1:5000 diluted horseradish peroxidase-labeled goat anti-mouse IgA were added. After washing the plate, 100 μL of OPD-citrate phosphate buffer solution was added, and after the substrate was developed, the reaction was terminated with 50 μL of 2 mol/L H 2 SO 4, and the A value at a wavelength of 490 nm was recorded by a microplate reader.

## Results and Discussion

SDS-PAGE of FTH, Glu, Glu-FTH protein purified Coomassie brilliant blue results are shown in Figures 1a, 1b, and 1c, respectively. The FTH protein was eluted by 200 mmol/L and 500 mmol/L imidazole. 60 mmol/L imidazole elutes Glu-FTH protein. The protein extraction and purification effects of this experiment are better, especially after FTH purification, the protein concentration and purity are higher. The extracted FTH and Glu-FTH are self-assembling nanoparticles. The protein was observed under transmission electron microscope. The diameters of Glu-FTH and FTH were about 20μm and 0.5μm, respectively. The size of the nanoparticles was uniform and the distribution was uniform. See Figure 2. 2.3 Anti-GluIgG levels in serum. The serum levels of anti-Glu IgG in each group were all at week 5. There was no significant difference between GluFTH, Glu, Glu-FTH+Poly(I:C), Glu+Poly(I:C) at week 5, but there was a statistical difference between them and FTH or PBS group (P <0.05), see Figure 3. In addition to the PBS and FTH groups, anti-GlusIgA levels in saliva of each group reached a high peak at week 5. At the 5th week, the level of anti-GlusIgA in the saliva of the Glu+Poly(I:C) group was significantly higher than that of the other groups.

**Figure 1.**
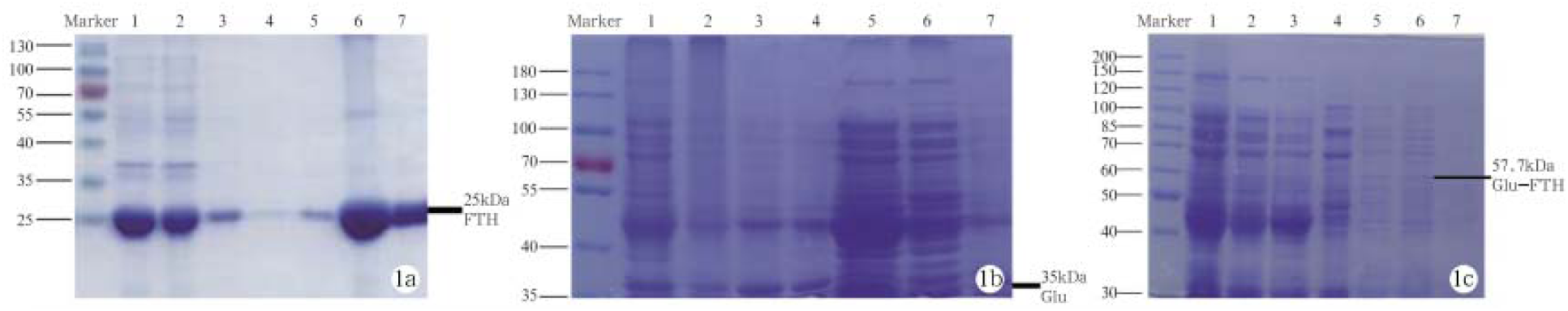
Protein content analysis by western blot

**Figure 2.**
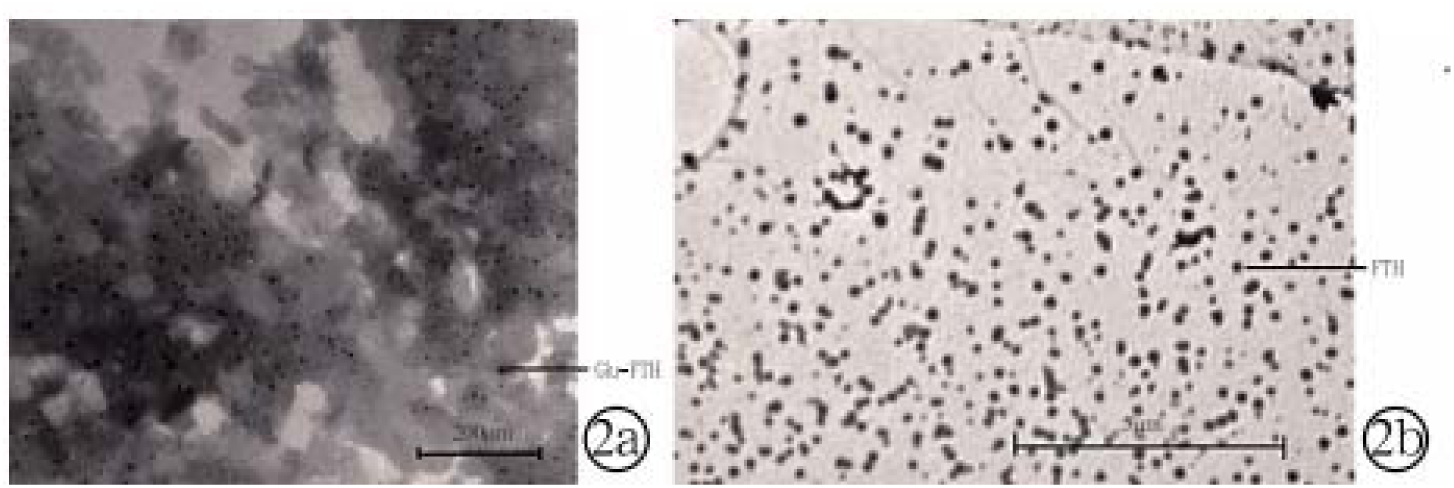
Electron microscopy analysis of self-assembled nanoparticles

**Figure 3.**
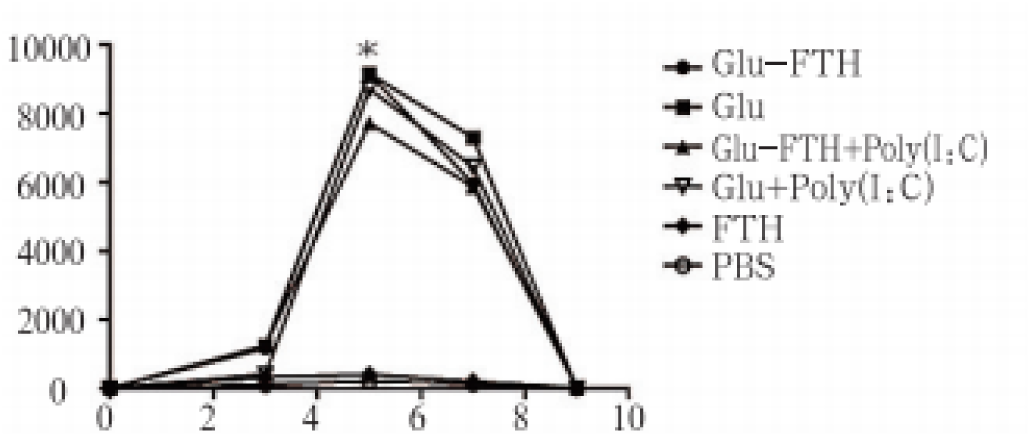
Antibody titer analysis

The types of adjuvants and the immune pathways can influence the type, intensity, and duration of immune responses [5]. Therefore, the selection of appropriate immunological adjuvants and immune pathways is important for improving the immune response of sputum-preventing vaccines. Scholars such as Steven GReed classify adjuvants into three categories: 1 non-immunogenic substances, which are used as antigen carriers to efficiently deliver more antigens to the immune system; 2 immunogenic substances that directly activate innate immunity Body (such as TLR); 3 both immunogenic and antigen can be presented to the immune system [6]. Poly(I:C) is a class 2 adjuvant with immunological double-stranded RNA molecules that enhance mucosal immunity through the TLR3 pathway and induce Th1-type immune responses [7]. Some scholars have combined Poly(I:C) with H1NI vaccine to achieve good immune effects [8]. The results of this study showed that the mucosal immune response produced by Poly(I:C) as an adjuvant combined with Glu immunized mice was stronger than that of Glu-immunized mice alone, and there was no significant difference in serum anti-GluIgG levels between the groups^1-14^. This confirms that Poly(I:C) is a better mucosal immune adjuvant that induces Th1-type immune responses [9].

Ferritin is a widely-preserved iron-storing protein that forms a cage structure composed of 24 subunits. The N-terminus exposed outside the cage structure can express and display exogenous proteins. Ferritin can be used as a carrier to fuse with a variety of antigens to form nanoparticles that present antigen to the immune system. There is insufficient evidence to suggest that FTH is immunogenic. Previous studies have used ferritin to present H1N1 protein subunit flu vaccine, which enhances the immunogenicity of HA antigens and induces the production of antibodies that neutralize multiple H1N1 subtypes [10]. Generally, the preparation of nanoparticles requires chemical treatment, and ferritin or its fusion protein can be self-assembled into nanoparticles, which reduces the damage of protein activity and facilitates the formation of uniform nanoparticles. In this study, a novel self-assembled nanoparticle anti-caries vaccine, Glu-FTH, was prepared by integrating the effective epitope of Streptococcus mutans Glu, which was screened by bioinformatics analysis, into the N-terminus of the ferritin cage structure by gene recombination. The FTH surface has a higher density of Glu epitopes, and Glu-FTH can be efficiently recognized and captured by antigen-presenting cells, thereby inducing a stronger immune response. In this experiment, Glu-FTH and Glu-FTH+poly (I:C) induced a certain amount of anti-GluIgG in serum at the 5th week, while Glu-FTH and Glu-FTH+poly(I: C) The concentration of anti-Glu-sIgA in the saliva of the group was lower. The size and physical and chemical properties of FTH may affect the immunological effects of Glu-FTH. Studies have shown that the size of the antigen carrier can affect the effect of the vaccine, however, the optimal size is still controversial; the physical and chemical properties of the antigen carrier can also affect the circulation of the antigen in the lymphatic system or the antigen presentation by the antigen presenting cells, thereby affecting the vaccine. Effect [11]. In addition, the results of SDSPAGE showed that there was still some hybrid protein (Glu-FTH) after purification of protein Glu-FTH. This heterologous protein may affect the immune effect of the vaccine. Our group will adjust the protein purification method to obtain higher purity. Protein. However, the ongoing experiments in this group (data not yet published) indicate that Glu-FTH has a significantly higher anti-PAcIgA level in saliva than Glu-immunized mice. It is suggested that saliva produced by Glu-FTH immunization has cross-protective effect on PAc, which may be due to the similar structure between Glu and PAc, suggesting that PAc-FTH (PAc and FTH fusion protein) may induce higher concentrations. Anti-PAcsIgA in saliva.

**Figure 4.**
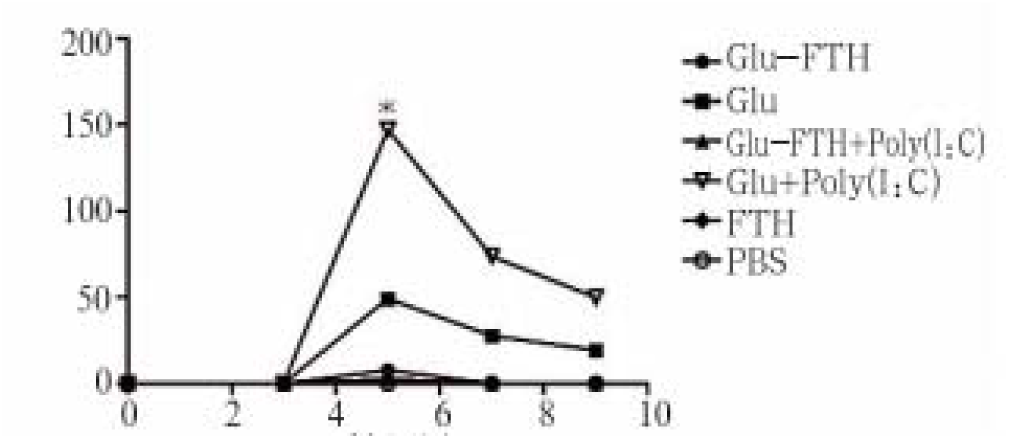
Antibody titer IgA analysis.

## Conclusion

In this report, we demonstrated that ferritin as an antigen-presenting vector also has a good application prospect. Nano-vaccine mucosal immune antigen presentation systems include virions, polyester microspheres, liposomes, and the like. The antigen can adhere to or be encapsulated on the surface of the antigen presenting system, and the presentation system can enhance antigen presentation, slow down antigen release, and thereby enhance antigen-specific immune responses. Moreover, the antigen presenting system has good biological properties such as biodegradability and biocompatibility. The US FDA has identified PLA and PLGA nanoparticles as molecular materials that can be used in humans and has been widely used as a drug delivery vehicle. Therefore, future efforts should focus on transforming this platform technology to the clinical settings.

## References

1. Laird, P. W.; Zijderveld, A.; Linders, K.; Rudnicki, M. A.; Jaenisch, R.; Berns, A., Simplified mammalian DNA isolation procedure. Nucleic acids research 1991, 19 (15), 4293.

2. Boussif, O.; Lezoualc’h, F.; Zanta, M. A.; Mergny, M. D.; Scherman, D.; Demeneix, B.; Behr, J.-P., A versatile vector for gene and oligonucleotide transfer into cells in culture and in vivo: polyethylenimine. Proceedings of the National Academy of Sciences 1995, 92 (16), 7297–7301.

3. Kenworthy, A. K.; Hristova, K.; Needham, D.; McIntosh, T. J., Range and magnitude of the steric pressure between bilayers containing phospholipids with covalently attached poly (ethylene glycol). Biophysical journal 1995, 68 (5), 1921–1936.

4. Krieg, A. M.; Yi, A.-K.; Matson, S.; Waldschmidt, T. J.; Bishop, G. A.; Teasdale, R.; Koretzky, G. A.; Klinman, D. M., CpG motifs in bacterial DNA trigger direct B-cell activation. Nature 1995, 374 (6522), 546.

5. Yu, Y.-C.; Roontga, V.; Daragan, V. A.; Mayo, K. H.; Tirrell, M.; Fields, G. B., Structure and dynamics of peptide-amphiphiles incorporating triple-helical proteinlike molecular architecture. Biochemistry 1999, 38 (5), 1659–1668.

6. Smith, J. D.; Cardwell, L. N.; Porciani, D.; Nguyen, J. A.; Zhang, R.; Gallazzi, F.; Tata, R. R.; Burke, D. H.; Daniels, M. A.; Ulery, B. D., Aptamer-displaying peptide amphiphile micelles as a cell-targeted delivery vehicle of peptide cargoes. Physical biology 2018, 15 (6), 065006.

7. Zhang, R.; Billingsley, M. M.; Mitchell, M. J., Biomaterials for vaccine-based cancer immunotherapy. Journal of Controlled Release 2018.

8. Zhang, R.; Kramer, J. S.; Smith, J. D.; Allen, B. N.; Leeper, C. N.; Li, X.; Morton, L. D.; Gallazzi, F.; Ulery, B. D., Vaccine Adjuvant Incorporation Strategy Dictates Peptide Amphiphile Micelle Immunostimulatory Capacity. The AAPS journal 2018, 20 (4), 73.

9. Zhang, R.; Leeper, C. N.; Wang, X.; White, T. A.; Ulery, B. D., Immunomodulatory vasoactive intestinal peptide amphiphile micelles. Biomaterials science 2018.

10. Zhang, R.; Leeper, C. N.; Wang, X.; White, T. A.; Ulery, B. D., Immunomodulatory vasoactive intestinal peptide amphiphile micelles. Biomaterials science 2018, 6 (7), 1717–1722.

11. Zhang, R.; Morton, L. D.; Smith, J. D.; Gallazzi, F.; White, T. A.; Ulery, B. D., Instructive Design of Triblock Peptide Amphiphiles for Structurally Complex Micelle Fabrication. ACS Biomaterials Science & Engineering 2018.

12. Zhang, R.; Smith, J. D.; Allen, B. N.; Kramer, J. S.; Schauflinger, M.; Ulery, B. D., Peptide Amphiphile Micelle Vaccine Size and Charge Influence the Host Antibody Response. ACS Biomaterials Science & Engineering 2018.

13. Zhang, R.; Smith, J. D.; Allen, B. N.; Kramer, J. S.; Schauflinger, M.; Ulery, B. D., Peptide Amphiphile Micelle Vaccine Size and Charge Influence the Host Antibody Response. ACS Biomaterials Science & Engineering 2018, 4 (7), 2463–2472.

14. Zhang, R.; Ulery, B. D., Synthetic vaccine characterization and design. Journal of Bionanoscience 2018, 12 (1), 1–11.

